# Optogenetic control of mechanotransduction based on light-induced homodimerization of talin

**DOI:** 10.1101/2025.04.17.649301

**Authors:** Ryosuke Nishimura, Samuel F. H. Barnett, Kashish Jain, Zengxin Huang, Benjamin T. Goult, Pakorn Kanchanawong

## Abstract

Integrin-based cell adhesions (IACs) serve as primary sites where piconewton-scale actomyosin-generated mechanical forces are transmitted to the extracellular matrix (ECM), generating traction forces that drive cell-ECM responses including adhesion, migration, and mechano-signaling. Talin, a large (270 kDa) cytosolic adaptor protein, is the principal force-transmission protein in integrin-based adhesions, containing multiple mechanosensitive domains and protein-protein interaction sites that orchestrate molecular events in mechanosensing. As a highly modular multi-domain protein, talin has been identified as an effective target for chemogenetic and optogenetic manipulation of integrin-based mechanotransduction. However, a key limitation of previous approaches is the reliance on heterodimerization modules to control talin function, requiring the expression of two modified talin fragments. In practice, achieving precise expression levels in such a 2-component approach can be challenging, particularly when combined with other genetic tools. Since talin naturally contains a C-terminal dimerization domain that forms part of its actin-binding site, we reasoned that the molecularly engineered talin with a C-terminal optically-controlled homodimerizer could enable single-component optogenetic control of mechanotransduction. This approach would facilitate multiplexing with other molecular perturbations or experimental techniques. Here, we describe an opto-homodimerizable talin based on the pdDronpa1.2 optogenetic module, which enables optogenetic control of talin by a single construct. We demonstrate that light-induced talin dimerization promotes talin recruitment to IACs, adhesion formation, actin retrograde flow engagement, and downstream mechanotransduction signaling. Conversely, light-induced talin monomerization rapidly disassembles focal adhesions, disrupts talin-actin linkages, and accelerates actin retrograde flow, underscoring the critical roles of talin dimerization. Furthermore, our single-construct design allows facile multiplexing of optogenetic modulation of integrin-mediated mechanotransduction with super-resolution single-molecule tracking, revealing the essential role of talin dimerization for integrin α_v_β5 engagement.

## Introduction

Integrin-based adhesions complexes (IACs) are multi-protein complexes that mediate cellular attachment to the extracellular matrix (ECM) (Kanchanawong and Calderwood, 2022). In adherent cells, integrin-based cell adhesions typically originate as diffraction-limited puncta, termed nascent adhesions, which undergo mechanosensitive maturation, progressing through distinct stages: focal complexes, focal adhesions (FAs), and fibrillar adhesions (Gardel et al., 2010). Heterodimeric αβ integrin transmembrane receptors physically couple to the actin cytoskeleton through a variety of cytosolic proteins, foremost of which is the talin family of adaptor proteins (Goult et al., 2018; Sun et al., 2019b). Talins bind the integrin β-cytoplasmic domain, stabilizing the activated conformation of integrin while directly and indirectly linking to actin filaments (Zhang et al., 2008). Notably, talin has been shown to transmit mechanical forces of up to ∼10 pN, which is critical not only for adhesion maturation but also mechanotransduction via associated signaling pathways (Austen et al., 2015; Kumar et al., 2016; Ringer et al., 2017; Zhou et al., 2017a). While the biochemical and biophysical properties of talin have been extensively characterized down to the structural level (Biertümpfel et al., 2025; Dedden et al., 2019; Goult et al., 2013a; Goult et al., 2013b; Yao et al., 2014), numerous aspects of how its mechanotransduction functions are dynamically regulated during cellular activities such as directed migration remain less well understood.

Structurally, talins are large (∼270 kDa), highly modular proteins (Yao et al., 2016; Yu et al., 2020) consisting of an N-terminal FERM domain that binds to integrin and a C-terminal rod domain that contains 13 alpha-helical bundles (R1-R13) (Goult et al., 2013b). The talin rod features multiple direct and indirect actin-binding sites (Critchley, 2009; Goult et al., 2013b; Haining et al., 2016; Yao et al., 2016). The actin-binding site 2 (ABS2), spanning R4-R8, has been associated with high-tension connection to F-actin in mature FAs (Kumar et al., 2016; Ringer et al., 2017). The C-terminal actin-binding site 3 (ABS3) consists of the 5-helical R13 domain and the single-helix C-terminal dimerization domain (DD) (Biertümpfel et al., 2025). The R13 domain is implicated in talin engagement with dendritic F-actin networks in lamellipodia (Driscoll et al., 2020), while the DD contributes to ABS3 actin binding and mediates talin dimerization (Biertümpfel et al., 2025; Gingras et al., 2008; Goult et al., 2013a). High-resolution CryoEM structures of autoinhibited talin revealed a monomeric conformation (Dedden et al., 2019), while dimeric talin has also been observed in an autoinhibited stacked torus configuration (Goult et al., 2013a). Dimerization at the very C-terminus confers bivalent integrin activation and cross-linking capability, which has been suggested to enhance integrin activation potency (Lu et al., 2022). A recent study further implicated talin dimerization in the incorporation of activated but unliganded integrin into adhesion complexes, as well as in mechanosensitivity to nanoscale ligand geometry (Changede et al., 2019). The dimerization state of talin thus has significant mechanobiological implications. Therefore, the ability to control the dimerization state of talin should provide a powerful tool for manipulating mechanobiological processes and dissecting mechanobiological mechanisms.

The recent development of conditionally dimerizable modules, using chemogenetic or optogenetic approaches, has enabled the control of protein-protein interactions with high spatiotemporal precision (Dagliyan and Hahn, 2019; Klewer and Wu, 2019; Oakes et al., 2017; Seong and Lin, 2021; Wittmann et al., 2020). As a highly modular protein, talin can be intensively modified through truncation or insertion to investigate various mechanobiological functions (Atherton et al., 2015; Austen et al., 2015; Driscoll et al., 2020; Liu et al., 2015b; Rahikainen et al., 2019). Recently, we and others have developed a toolbox of chemogenetically and optogenetically controlled talins to dissect talin-mediated force transmission in integrin-based cell adhesions (Sadhanasatish et al., 2023; Wang et al., 2019; Yu et al., 2020). However, many of these systems rely on heterodimerization modules, requiring the expression of multiple components at comparable level in cells. This complexity presents a major practical limitation, making it challenging to multiplex chemo/optogenetic tools with microscopy techniques that require additional genetically encoded probes. Achieving precise expression levels of multiple constructs within a single cell remains difficult. Although multi-cistronic expression cassettes (e.g. internal ribosome entry site (IRES) or self-cleavable 2A peptides) offer potential solutions, they suffer from drawbacks such as low expression efficiency or self-cleavage failure (Bochkov and Palmenberg, 2006; Kim et al., 2011). To address these limitations and expand the utility of talin-based mechano-optogenetics, we implemented a reversible homodimerizing optogenetic module, pdDronpa1.2 (Zhou et al., 2017b) to achieve light-inducible control of talin monomer/dimer transition. Due to the homodimerization property of pdDronpa1.2, our system requires only a single construct to control integrin-based cell adhesions optogenetically. Upon 488-nm illumination adhesion disassembly occurs with high spatiotemporal specificity. Our optodimerizable-talin reveals that talin dimerization—whether native or artificially induced—is essential for talin recruitment, adhesion formation, and actin cytoskeleton engagement. Furthermore, we demonstrate the multiplexing of optogenetic adhesion control with single-molecule analysis, providing insights into how adhesions regulate membrane-associated protein dynamics.

## Results

### Modulating talin function by a mechanically robust reversible optical dimerizer

The fluorescent protein pdDronpa1.2 has recently been demonstrated to form a homodimer that can be reversibly dissociated by 488 nm illumination, while re-dimerization is promoted by 405 nm illumination (Zhou et al., 2017b). Furthermore, pdDronpa1.2 (henceforth referred to as Dronpa for short) undergoes a photochromic shift in conjunction with a monomer/dimer transition. In the dimeric state, Dronpa is green-fluorescent (488 nm excitation channel), while the monomeric state is largely non-fluorescent, allowing the dimerization state to be readily monitored. Earlier characterization by Atomic Force Microscopy has shown that Dronpa is capable of sustaining a high level of mechanical force up to ∼80 pN *in vitro* (Jöhr et al., 2019), well beyond the previously measured magnitude of intramolecular tension in talin (Austen et al., 2015; Kumar et al., 2016). Since dimer rupture force may vary as a function of the loading rate, we first sought to ascertain whether Dronpa is capable of supporting physiological force as sustained by talin in living cells. As shown in **Fig. 1a**, we generated a pair of Dronpa-fusion constructs with the first containing an N-terminal mApple fused to residue 1–1973 (FERM and R1-R10 domains) of mouse talin1 and a C-terminal Dronpa, and the second containing an N-terminal Dronpa fusion to residue 1974–2541 (R11–R13 and DD domains) and a C-terminal mIFP (Yu et al., 2015). Upon Dronpa homodimerization, if these constructs are capable of sustaining pN force, they are expected to restore mechanical connection between integrin and actin cytoskeleton through talin, thereby rescuing talin-deficient phenotypes and enabling FA formations and cell migration. Conversely, Dronpa monomerization is expected to sever the mechanical connection between integrin and actin, leading to FA disassembly and cell retraction (**Fig. 1b**).

**Figure 1.**
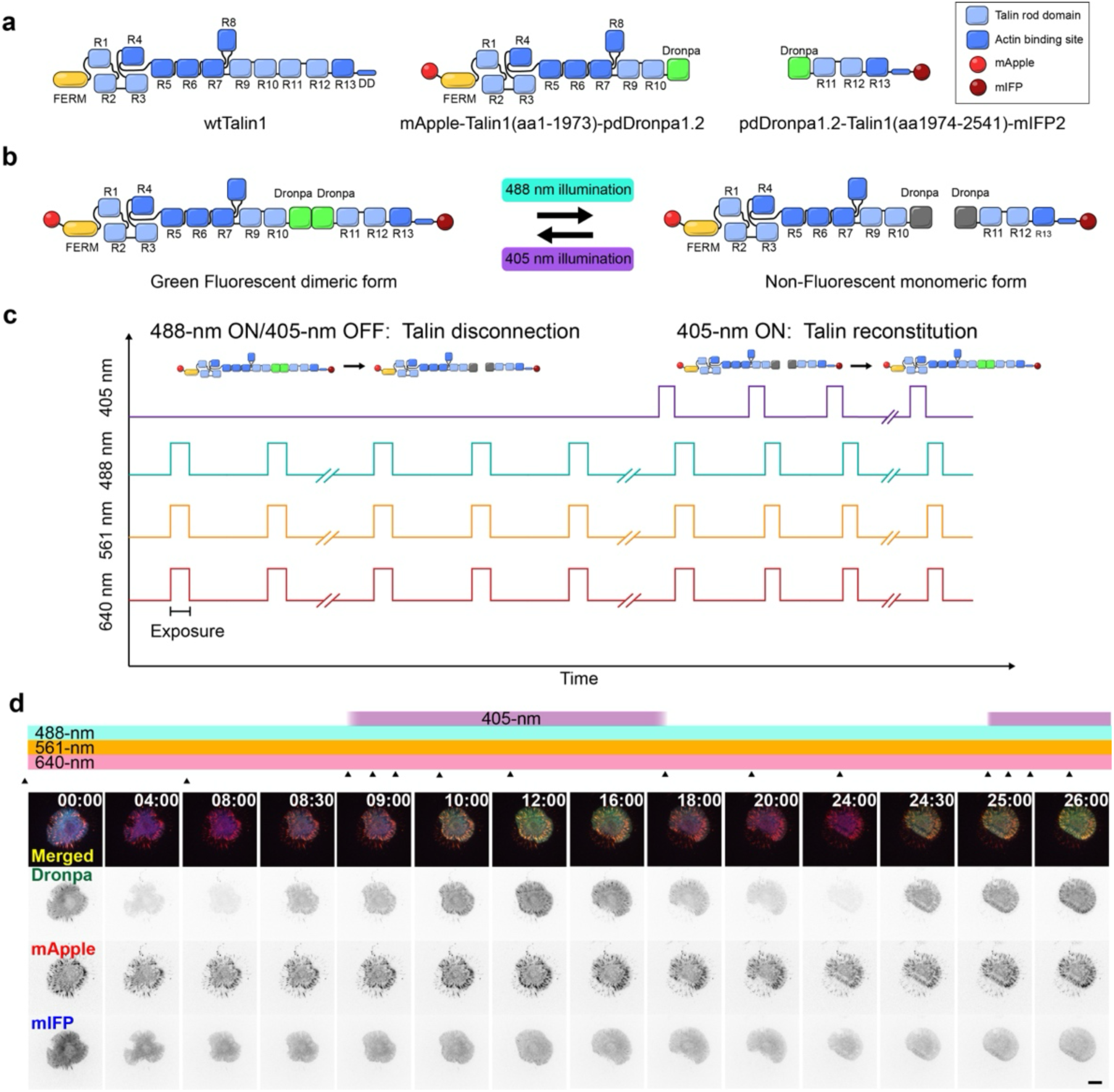
Multi-spectral optogenetic control of talin mechanical connection by Dronpa. **a**) Schematic diagram of (left) full-length talin1 (wtTalin1; residues 1–2541), (center) N-terminal optogenetic element containing truncated murine talin1 (residues 1–1973) and C-terminally fused Dronpa, (right) C-terminal optogenetic element containing Dronpa fused N-terminally to talin1 fragment (residues 1974–2541). As the optogenetic elements are respectively fused by mApple and mIFP, thisallows for fluorescent monitoring in the red and far-red channels. **b**) Schematic diagram for Dronpa-mediated reversible reconstitution of full-length talin. (Left) Dronpa dimerization (green fluorescent) forms a mechanically intact talin able to mediate integrin-actin connection in FAs. (Right) 488-nm illumination leads to a reversible photoconversion of Dronpa to monomeric and non-fluorescent form, leading to the disruption of talin mechanical connection. 405-nm illumination of monomeric Dronpa, in turn, photoconverts Dronpa back to the dimeric and green-fluorescent form, restoring the mechanical connection. **c)** Reversible control of FA assembly/disassembly by Dronpa dimerization of talin fragments. **d)** Selected frames from a time-lapse video (**Movie 1**) showing alternating phases of FA maturation and cell protrusion (during 405-nm illumination) and FA disassembly and cell retraction (during 488-nm illumination). Scale bar, 10 µm.

When co-transfected into *talin1^-/-^talin2^-/-^*fibroblasts (TKO) cells (Austen et al., 2015), which are weakly adherent and unable to form FAs, we found that these constructs rescue and localize to FAs (**Fig. 1d**) and that dimerization can be reversibly modulated by 405-nm and 488-nm illumination (**Fig. 1c,d**, **Movie 1**). Upon illumination by 488-nm, the dimer form of Dronpa can be observed by the green fluorescence. The concomitant robust FA maturation and efficient cell spreading (**Fig.1d** leftmost panel) indicates that the Dronpa dimer can form an effective mechanical connection that reconstitutes the functionality of full-length talin. Following several minutes of 488-nm illumination, a reduction in green fluorescence is observed which can be interpreted as the dissociation of dimeric Dronpa in conjunction with photoconversion into the non-fluorescent form (**Fig. 1b**). Consistent with this, upon illumination at 405-nm we observed a rapid increase of green fluorescence in FAs, which can be interpreted as the re-formation of fully-connected talin in conjunction with FA maturation and increased cell spreading. By cycling the 405-nm and then 488-nm illuminations, we were able to observe multiple cycles of FA formation and disassembly in conjunction with the appearance and disappearance of green fluorescence (**Fig. 1d**, **Movie 1**). Based on previous measurements of force magnitude in talin (Ringer et al., 2017), our results indicate that the pdDronpa1.2 dimer is mechanically robust and thus suitable as a reversible, multi-spectra dimerizer for investigating mechanotransduction processes in live cell contexts.

### Optical control of talin1 dimerization

The talin dimerization domain (DD) consists of 47 amino acids (residues 2496–2529) at the very C-terminus of talin. Earlier biochemical analysis has shown that the DD domain can form an antiparallel coiled-coil (Gingras et al., 2008), thus the DD domain is thought to mediate the formation of the talin-dimer (Goult et al., 2013a). In previous studies, talin truncations that eliminate the DD domain generally abrogate FA formation, except when additional talin-actin crosslinking is also provided, such as by co-expression with constitutively active vinculin (Atherton et al., 2015). To test whether DD is essential for talin activity within FAs, we generated a deletion construct of mouse talin1 lacking the DD domain (ΔDD: 1–2493) tagged at the N-terminus by mIFP for visualization (**Fig. 2a**). In weakly adherent TKO cells, the expression of wild-type talin1 is sufficient to rescue cell spreading and FA formation (**Fig. 2d,f**). In contrast, upon the expression of ΔDD in TKO cells, we observed that while cells are able to spread on the fibronectin-coated surface and form actin-rich lamellipodia (due to the activity of talin head domain in ΔDD), the spreading areas were small while the cells remained highly circular and unable to polarize (**Fig. 2d,f**). We also observed that FAs, defined as micron-scale protein plaques, are absent in virtually all ΔDD cells, with occasional indistinct adhesions observed in a few cells reminiscent of earlier observations in talin mutants (Azizi et al., 2021). Morphologically, the ΔDD cells were similar to TKO cells either expressing the talin1 head domain or treated with Mn^2+^ (**Fig. S1**), which can activate integrins but are unable to connect integrin to the actin cytoskeleton. Interestingly, even when co-expressed with wt-talin1, ΔDD failed to localize to adhesions and remained cytoplasmic (**Fig. S2a**). Our observations thus corroborate that the DD domain of talin is required for FA formation but also for talin localization to adhesions.

**Figure 2.**
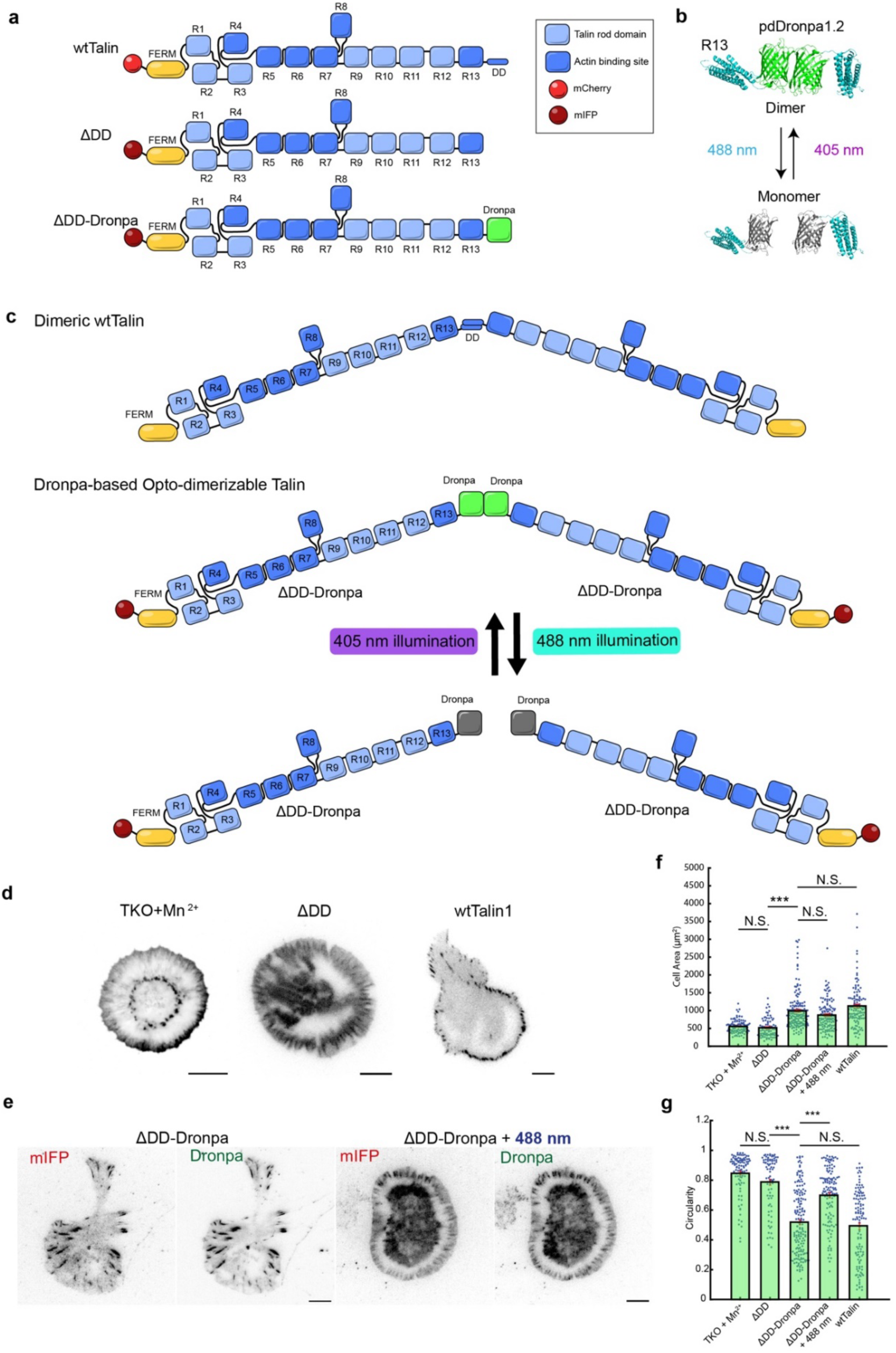
Opto-dimerizable talin1. **a)** Schematic diagram of talin N-terminally tagged with mCherry (mCh-wtTalin), mIFP-tagged ΔDD and ΔDD-Dronpa. wtTalin is full-length talin1 (residues 1–2541), ΔDD is truncated before the dimerization helix (1–2493), ΔDD-Dronpa is ΔDD followed by a short linker and Dronpa. **b)** Structural model of Dronpa-mediated Talin dimerization. Dronpa (green = dimer/fluorescent, grey = monomer/dark) attached to Talin-R13 domain (blue) in both the dimeric and monomeric forms. The dimer-monomer status can be controlled with 488 nm and 405 nm light. **c**) Schematic diagram for the dimeric form of talin and Dronpa-based opto-dimerizable-talin. **d)** Representative images of talin1/talin2 knockout (TKO) cells treated with Mn^2+^ and stained with Phalloidin-CF405, or transfected with ΔDD or wtTalin, plated on fibronectin-coated substrate. **e**) Representative images of TKO cells transfected with ΔDD-Dronpa in the dimeric state (left) and in the monomeric state induced by 3.6 W/cm^2^ 488-nm stimulation (right; 100-ms pulse every 30 seconds for 1 h). **f, g)** Quantification of cell spreading area (**f**) and cell circularity (**g**) (*n* = 158-328). ***: *p* < 5 x 10^-4^; N.S.: not significant; *P*-values and significance values were found with Kruskal-Wallis test with Tukey’s honest significance test. Scale bars, 10 µm.

To dynamically control talin dimerization, we next designed an opto-dimerizable construct, termed ΔDD-Dronpa, in which the Dronpa module replaced the DD domain, along with an N-terminal fusion of mIFP as a near-infrared fluorescent reporter (i.e. mIFP-linker-[mouse talin1 residue 1–2493]-linker-Dronpa, **Fig. 2a–c**). As shown in **Fig. 2e**, we observed that the expression of ΔDD-Dronpa in TKO cells rescued the formation of FAs, especially at the cell periphery, and resulted in a significant increase in cell spreading area and cell polarization compared to ΔDD (**Fig 2f,g**). Since dimeric Dronpa is green-fluorescent, while monomeric Dronpa is non-fluorescent (Zhou et al., 2017b), we next used Total Internal Reflection Fluorescence (TIRF) microscopy to monitor the monomerization and dimerization dynamics of ΔDD-Dronpa by observing 488 nm-channel fluorescence as an indicator for dimerization and 640 nm-channel fluorescence as an indicator for total protein. As shown in **Fig. 2e**, the high level of green fluorescence signal in FAs indicated that a significant proportion of ΔDD-Dronpa localizes to FAs in the dimeric form. Upon continued 488 nm illumination, Dronpa fluorescence became progressively attenuated **(Fig. S2b,c**), indicative of the transition from dimeric to monomeric forms. Concomitantly, monomerization is accompanied by a reduction in numbers of FAs and a decrease in cell polarization (**Fig. 2f,g**).

Next, we further characterized how the dimerization state of ΔDD-Dronpa influences the dynamics of cell spreading. We monitored the spreading of newly plated TKO cells expressing ΔDD-Dronpa under continuous 488-nm illumination at a power density of 0 (control), 1.3, or 3.6 W/cm^2^, with the pulse duration of 100 ms and 1 pulse per minute frequency (these illumination parameters were chosen to maintain Dronpa in the monomeric state while minimizing phototoxicity). We observed that cells exposed to 488-nm illumination retracted in a light-dependent manner whereas the control cells continued to increase the spreading area, as quantified in **Fig. 3a**. During mesenchymal cell motility, FAs undergo cycles of formation, maturation, and disassembly in a spatiotemporally coordinated fashion (Gardel et al., 2010; Kanchanawong and Calderwood, 2022; Parsons et al., 2010). Time-lapse TIRF imaging of the mIFP channel revealed that TKO cells expressing ΔDD-Dronpa were motile, with FAs undergoing formation, maturation, and disassembly as the cells migrate over fibronectin-coated substrates (**Fig 3b**, **Movie 2**).

**Figure 3.**
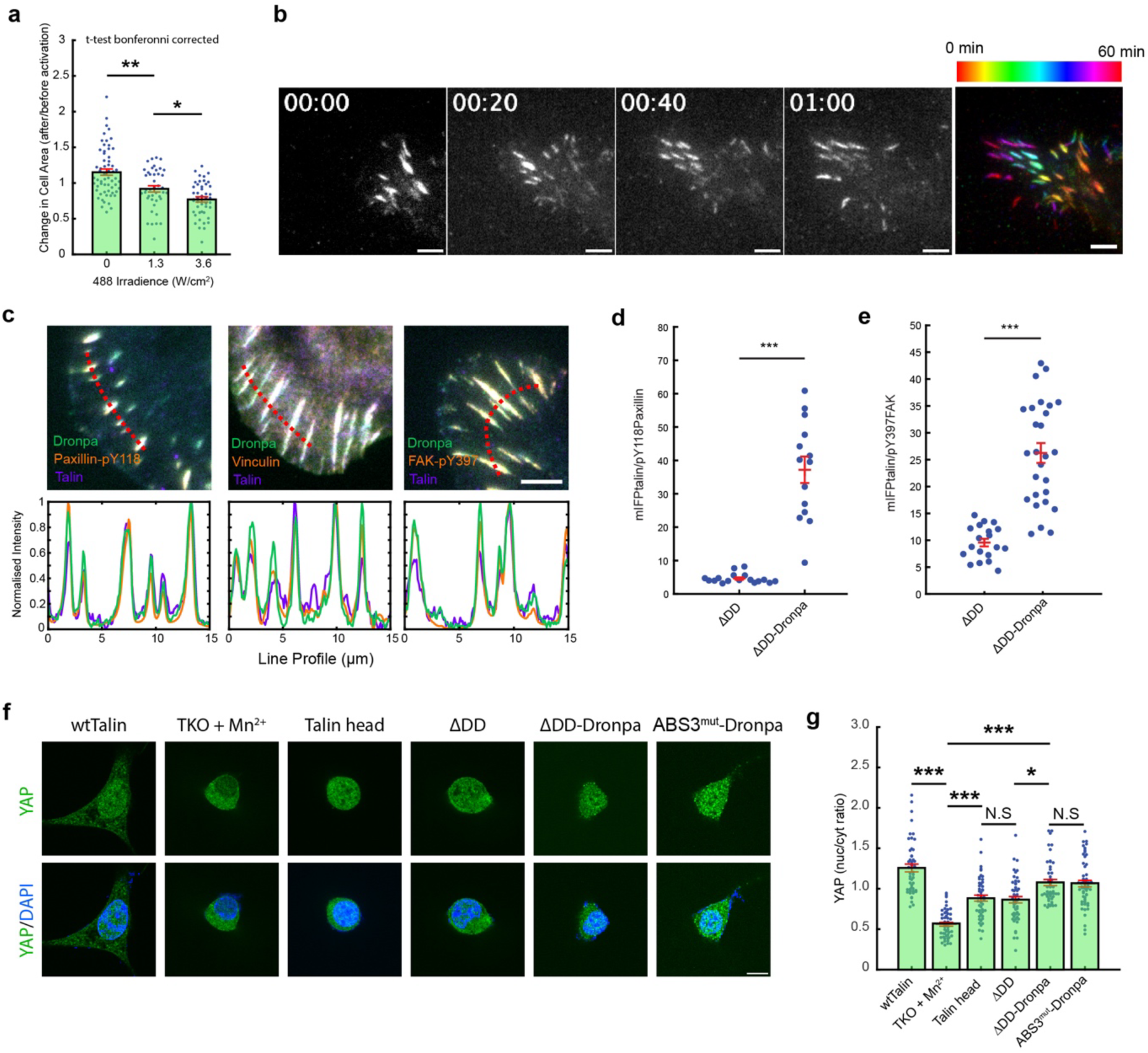
Characterization of FA maturation and dynamics mediated by opto-dimerizable talin1. **a)** Effects of monomerizing 488-nm illumination on the spread areas of TKO cells expressing ΔDD-Dronpa following 1 h of illumination at various intensities. **b)** Selected frames and temporal color-coded time-lapse image of ΔDD-Dronpa (mIFP channel) observed over 1 h demonstrating dynamics of FA turnover (*n* = 84–120; **Movie 2**). **c)** Merged immunofluorescence micrographs with immunostaining for paxillin-pY118, vinculin, and FAK-pY397 (orange) in TKO cells expressing ΔDD-Dronpa (green), with mIFP channel (purple) indicating total talin level. Line profiles (lower panels) correspond to red dotted lines on top panels. **d)** Comparison of activated paxillin signified by phosphotyrosine 118 between cells expressing Tln1ΔDD and Tln1ΔDD-Dronpa. **e)** Comparison of activated FAK as indicated by phosphotyrosine 397 between cells expressing Tln1ΔDD and Tln1ΔDD-Dronpa. **f)** Immunostaining for YAP shows that dimerization is important for talin-mediated YAP translocation to the nucleus. **g)** Quantification for the images shown in d, (*n* > 49 each condition). *: *p* < 5 x 10^-2^; **: *p* < 5 x 10^-3^; ***: *p* < 5 x 10^-4^; N.S.: not significant; *P*-values and significance values were found with Kruskal-Wallis test with Tukey’s honest significance test. Scale bars, 10 µm (**b),** 5 µm (**c,f**).

### FAs formed by ΔDD-Dronpa**-**talin support integrin-mediated mechanosignaling

Next, we sought to further characterize the ability of opto-dimerizable ΔDD-Dronpa-talin to recapitulate mechanosignalling functions of wild-type talin. From immunofluorescence microscopy of TKO cells expressing either ΔDD or ΔDD-Dronpa constructs, we observed prominent co-localization of vinculin with ΔDD-Dronpa (**Fig. 3c**) but not ΔDD (**Fig. S2d**), as expected. Markers of integrin-dependent signaling such as active (open conformation) integrin β1 (**Fig. S2e**), phospho-Paxillin (pY118) (Zaidel-Bar et al., 2007), which is phosphorylated by FAK and Src, and phospho-FAK (pY397) (Chen et al., 1996), were also observed to co-localize with ΔDD-Dronpa in FAs (**Fig. 3c, Fig. S2d**). Furthermore, when normalized to talin and compared to ΔDD, the levels of phospho-paxillin and phospho-FAK staining in ΔDD-Dronpa cells were markedly increased indicative of higher levels of activation when talin undergoes dimerization (**Fig. 3d,e**). Collectively, these data suggest that integrin-mediated mechanotransduction signaling can be recapitulated by ΔDD-Dronpa talin.

We also assessed whether ΔDD-Dronpa talin is able to recapitulate talin-mediated mechanotransduction at the transcriptional level by monitoring nuclear translocation of YAP, a canonical mechanosensitive transcription factor known to be responsive to talin-dependent mechanotransduction (Dupont et al., 2011; Elosegui-Artola et al., 2016). From immunofluorescence microscopy of endogenous YAP, we quantified the nucleus-to-cytoplasmic ratio of YAP intensity, as shown in **Fig. 3f,g**. TKO cells with re-expression of full-length talin1 exhibited high nuclear localization of YAP, indicative of robust downstream mechanosignalling. Since control TKO cells are weakly adherent, Mn^2+^ treatment was used to promote integrin activation and cell adhesion, while the lack of connection between integrin and actin cytoskeleton gave rise to low nuclear localization of YAP, as expected (**Fig. 3f,g**). We also observed that TKO cells expressing talin head domain or ΔDD-talin exhibit relatively low nuclear YAP compared to full-length talin expression. On the other hand, re-expression of ΔDD-Dronpa-talin was able to rescue YAP nuclear translocation up to 74% compared to full-length talin1. Interestingly, we also observed that ΔDD-Dronpa-talin with KVK/DDD mutation that disrupts ABS3 (ABS^mut^) (Atherton et al., 2015; Bate et al., 2012), is also able to rescue nuclear translocation to a similar extent as ΔDD-Dronpa-talin, suggesting that dimerization of talin may be an important determinant of YAP signaling.

### Talin dimerization is required for the engagement of molecular clutch to actin retrograde flow

As the presence of the DD domain in wildtype talin is essential for FA maturation, we next sought to assess its involvement in the engagement with actin retrograde flow. To simultaneously monitor the actin flow rate and adhesions dynamics, we co-expressed an F-actin probe, F-tractin-mCherry in conjunction with the talin constructs in TKO cells and performed time-lapse imaging by spinning-disk confocal microscopy. Kymograph analysis of cell protrusive edge as well as Particle Image Velocimetry (PIV) analysis of the F-tractin channel were then used to quantify actin retrograde flow. We observed that TKO cells expressing only F-tractin-mCherry or ΔDD exhibited a similarly rapid actin retrograde flow of ∼175 nm/s as quantified by PIV analysis **(Fig. 4a,b, Movies 3–6)**. In terms of the molecular clutch model, these high flow rates can be interpreted as being due to the lack of molecular clutch engagement between integrin and actin retrograde flow, either due to the absence of talin or the ineffective clutch function of ΔDD construct. Importantly, in ΔDD-Dronpa cells we observed that the actin retrograde flow becomes significantly reduced to ∼110 nm/s suggesting that ΔDD-Dronpa can engage and thus slow down actin retrograde flow significantly, although not to the same extent as wildtype talin GFP (∼70 nm/s) **Fig. 4a,b**. Notably, in ΔDD-Dronpa and wtTalin cells we also observed the formation of distinct actin bundling at the lamella-lamellipodia border (Burnette et al., 2014), while in ΔDD cells, this bundling is largely absent (**Movies 3–6**). Moreover, taking advantage of optically-induced dimerizability of ΔDD-Dronpa, we tested whether actin retrograde flow impedance depends on talin dimerization. As shown in **Fig. 4c, d** and **Movie 7,** in unstimulated ΔDD-Dronpa cells, we observed a lamellipodial actin retrograde flow of 110 nm/s. Upon 488-nm illumination for 300 s to monomerize talin, we observed the acceleration of actin retrograde flow to 206 nm/s. Subsequently, upon 405-nm illumination to re-dimerize talin, we observed that actin retrograde flow decelerated again to 154 nm/s. Consistent with this, upon 488-nm induced talin monomerization, the lamella-lamellipodia bundles appear to disassemble in conjunction with the reorganization of the lamellipodial actin (**Fig. S3**). Taken together, these results indicated that the dimerization state of talin plays an important role in regulating actin retrograde flow engagement and actin cytoskeletal organization in a manner consistent with the molecular clutch model. This further demonstrates how our opto-dimerizable talin can dynamically exert direct control of the FAs molecular clutch.

**Figure 4:**
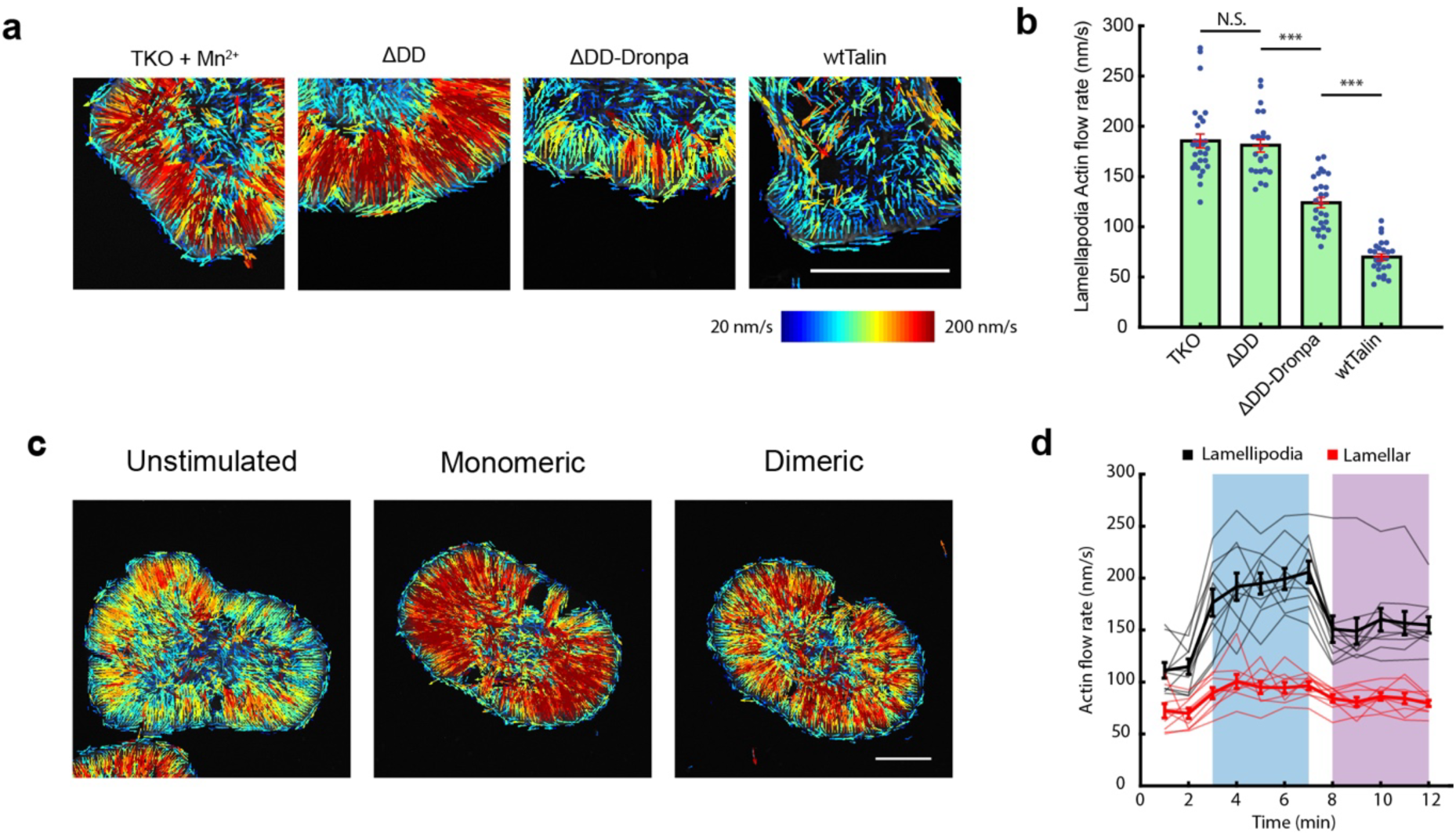
Talin dimerization controls engagement with the actin cytoskeleton. **a)** Particle Image Velocimetry (PIV) analysis of retrograde actin flow at the cell edge of TKO cells spread with manganese ions or transfected with ΔDD, ΔDD-Dronpa, or wtTalin (**Movies 3–6**). **b)** Quantification of PIV analysis in **a**). **c)** Single cell expressing ΔDD-Dronpa analyzed by PIV analysis in the unstimulated-dimer (left), monomeric (middle), and redimerized (right; **Movie 7**). **d)** Time-course of actin retrograde flows for cells stimulated with 488-nm light to monomerize talin, followed by 405-nm light to redimerize talin. Black represents actin flow in the lamellipodia whereas red represents actin flow in the lamellar region. Pale lines represent individual cell time-courses and the thick line is the average response. Scale bars, 10 µm.

### Spatiotemporally precise control of talin dimerization in FAs

Our results thus far showed that, in lieu of the DD domain, the Dronpa moiety in the opto-dimerizable talin appears to be capable of rescuing structural and localization properties of talin to recapitulate significant aspects of talin functions. These also implicate talin dimerization as being necessary for both the formation and the continuous maintenance of FAs, efficient cell spreading, and cell polarization. While these suggest that endogenous talin is likely in the dimeric form when localized to FAs, since talin could also be present in monomeric form (Dedden et al., 2019), we next investigate the effects of talin monomerization in FA organization and dynamics by making use of the spatiotemporal precision offered by ΔDD-Dronpa. First, to dissect the contribution of dimerization to FA maturation and talin recruitment, we applied cyclic illumination that alternates between 405-nm (dimerizing) and 488-nm (monomerizing) excitation every six minutes. Each cycle contained a pulse train consisting of 200 ms pulses of either 488-nm or 405-nm light with an interval of 5 seconds as shown in **Fig. 5a–c** (see also **Movie 8**). Consistent with earlier observation, during the 405-nm illumination phase, we observed a robust growth in the adhesions area and a significant increase in talin recruitment as determined by the N-terminal mIFP signal. In contrast, during 488-nm illumination the abundance of adhesion-localized talin either reduced or stagnated, though the intensity decrease was generally uniform over the adhesions (**Fig. 5a–c**). These results suggest that the dimeric form of talin can be readily incorporated into FAs to contribute to maturation. On the other hand, acute disruption of the dimer appears to result in the rapid dissociation of talin from FAs, hence a relatively isometric reduction of talin abundance. We also observed that cyclic monomerization/dimerization illumination visibly causes the cell body to alternately shift away from or toward the leading edge of FAs in synchrony with the illumination cycles, suggesting that the adhesion-cytoskeleton linkage and actomyosin contractility may also be modulated in conjunction with talin dimerization (see **Movie 8**).

**Figure 5.**
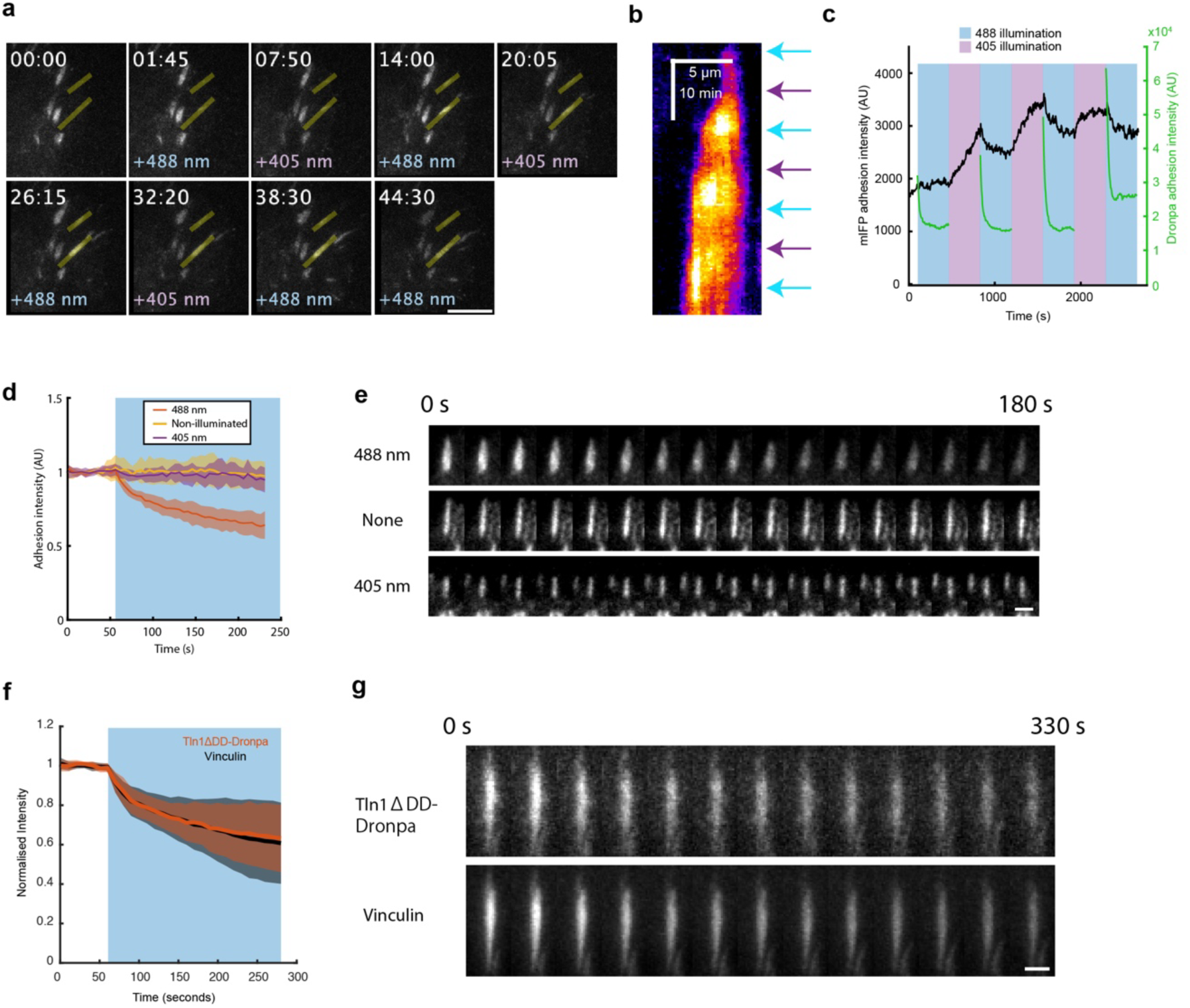
Spatiotemporal modulation of talin dimerization **a)** Still images of TKO cells expressing ΔDD-Dronpa, during cyclic illumination at specified ROI (Yellow overlay). **b)** Kymograph of a FA exposed to cyclic 488/405-nm light (indicated by arrows, cyan is 488-nm, purple is 405-nm). **c)** Intensity profile of adhesion in **b)** showing that mIFP intensity in the adhesion (black) correlates with illumination wavelength, also shown is the Dronpa fluorescence decay (green), (**Movie 8)**. **d)** mIFP intensity of individually targeted adhesions illuminated with either 488-nm light, 405-nm light, or no stimulation. **e)** mIFP intensity of example adhesions from **d)** (**Movie 9)**. **f**) Levels of Vinculin drop in conjunction with levels of talin as blue light monomerizes ΔDD-Dronpa. **g)** Dronpa and mCherry-Vinculin signal of example adhesions from **f)** (**Movie 10)**. Scale bars, 10 µm (**a**), 2 µm (**e, g**).

We next investigated the effects of talin dimerization at the single-FA level. Using a fluorescence recovery after photobleaching (FRAP) module to selectively illuminate individual FAs with 488-nm light, we observed a significant reduction of the mIFP intensity within the illuminated region of interest, decreasing by ∼40% over a 3-minute period. In contrast, nearby non-illuminated adhesions remained unaffected and maintained their mIFP intensity (**Fig. 5d,e; Movie 9**) over the same period. We interpreted this as being due to the dissociation of monomeric talin upon 488-nm-induced monomerization. Consistent with this, in cells co-transfected with ΔDD-Dronpa with mCherry-Vinculin, we observed that upon 488-nm illumination of individual FAs, a synchronous loss of vinculin is observed that mirrored talin vacating the adhesion (**Fig. 5f,g; Movie 10**). Interestingly, upon selective illumination of FAs using 405-nm light, we observed that FA area and talin abundance remained largely unchanged. These results suggest that talin may be fully dimeric inside adhesions, hence additional 405-nm stimulation does not increase dimeric talin further.

### Optodimerizable-talin enables multiplexing of optogenetics and single-molecule fluorescence imaging

Single-molecule-based microscopy techniques have been instrumental in elucidating molecular-scale organization in a broad range of cellular structures (Barnett and Kanchanawong, 2018; Bertocchi et al., 2013; Xia and Kanchanawong, 2017), including unveiling the spatial heterogeneity of FA protein dynamics within nanoscale compartments in FAs or between different plasma membrane regions (Fujiwara et al., 2023a; Fujiwara et al., 2023b; Huang and Kanchanawong, 2023; Liu et al., 2021; Massou et al., 2020; Rossier et al., 2012). For instance, the mobility and density of integrins are found to be modulated locally relative to protrusion onset (Jaqaman et al., 2016), while individual integrins were observed to undergo repeated cycles of immobilization and diffusion, traversing adjacent FAs within minutes, with increased immobilization toward the distal FA tips presumably due to higher traction (Tsunoyama et al., 2018). These observations suggest that membrane and protein dynamics at the single-molecule level are likely interdependent with global cellular behavior, but the ability to probe such process in conjunction with precise spatiotemporal perturbation has been less well developed. To demonstrate the utility of Dronpa-based optodimerizable-talin1 in multiplexing optogenetic control with single-molecule analysis, we therefore performed single-particle tracking (SPT) analysis to evaluate the dynamics of the RGD-binding integrins α_V_β3 and α_V_β5 in response to FA perturbation. In particular, while α_V_β3 are widely known to be early components of FAs and whose activations are regulated by talin (Kanchanawong and Calderwood, 2022), for α_V_β5, in addition to FA localization, localization in reticular adhesions (RAs) in a talin1 and actin-independent manner has also been reported (Lock et al., 2018).

As control, we first performed SPT analysis in TKO cells treated with Mn^2+^ to activate integrin and enhance cell spreading. Integrin β3 or integrin β5 C-terminally fused to HaloTag (β3-Halo and β5-Halo) were then stably expressed, whereby heterodimerization with endogenous integrin α_v_ is expected to form integrin α_V_β3 or α_V_β5, respectively. In conjunction, the plasma membrane probe CAAX-HaloTag was used to probe plasma membrane dynamics. The fluorophores JF549 or JF646 were used to label HaloTag (**Fig. S4**) and to enable simultaneous imaging with the Dronpa fluorescence channel when applicable. Single-molecule time-lapse movies were collected using 640-nm TIRF illumination, with examples of single-molecule movies in **Movies 11–19**. Single-molecule coordinates were localized and linked trajectories longer than 25 frames were used to calculate diffusion coefficients as described previously (Rossier et al., 2012) (Fig. 6a). Categorization between mobile and immobile trajectories was performed using the diffusion coefficient threshold of 0.01 µm/s^2^ similar to in previous studies (Rossier et al., 2012). As expected, we observed that the CAAX-HaloTag membrane probes in control TKO cells (Mn^2+^) are highly mobile (Fig. 6a). The histogram plot of diffusion coefficients revealed a unimodal distribution peaking at D ∼0.1 µm/s^2^ (Fig. 6b). Classification of CAAX trajectories revealed that the majority of trajectories are diffusive, while immobilized particles comprise only 25% (Fig. 6c). We observed that integrin β3 trajectories are largely mobile similar to CAAX (Fig. 6a), likely due to the absence of talin. In contrast, for integrin β5, a majority of the trajectories are found to be immobile (72%), suggestive of their ability to become activated independently of talin as described earlier (Lock et al., 2018). Next we performed SPT analysis in TKO cells rescued by stable expression of talin1-mEmerald to enable FA formation. Using the talin1-mEmerald channel to segment FA regions, SPT analysis reveals that the CAAX membrane probes remains highly mobile both inside and outside of FAs, while for both integrin β3 and β5 a significant increase in immobile trajectories are observed inside FAs as expected (Fig. 6d**–f**).

**Figure 6:**
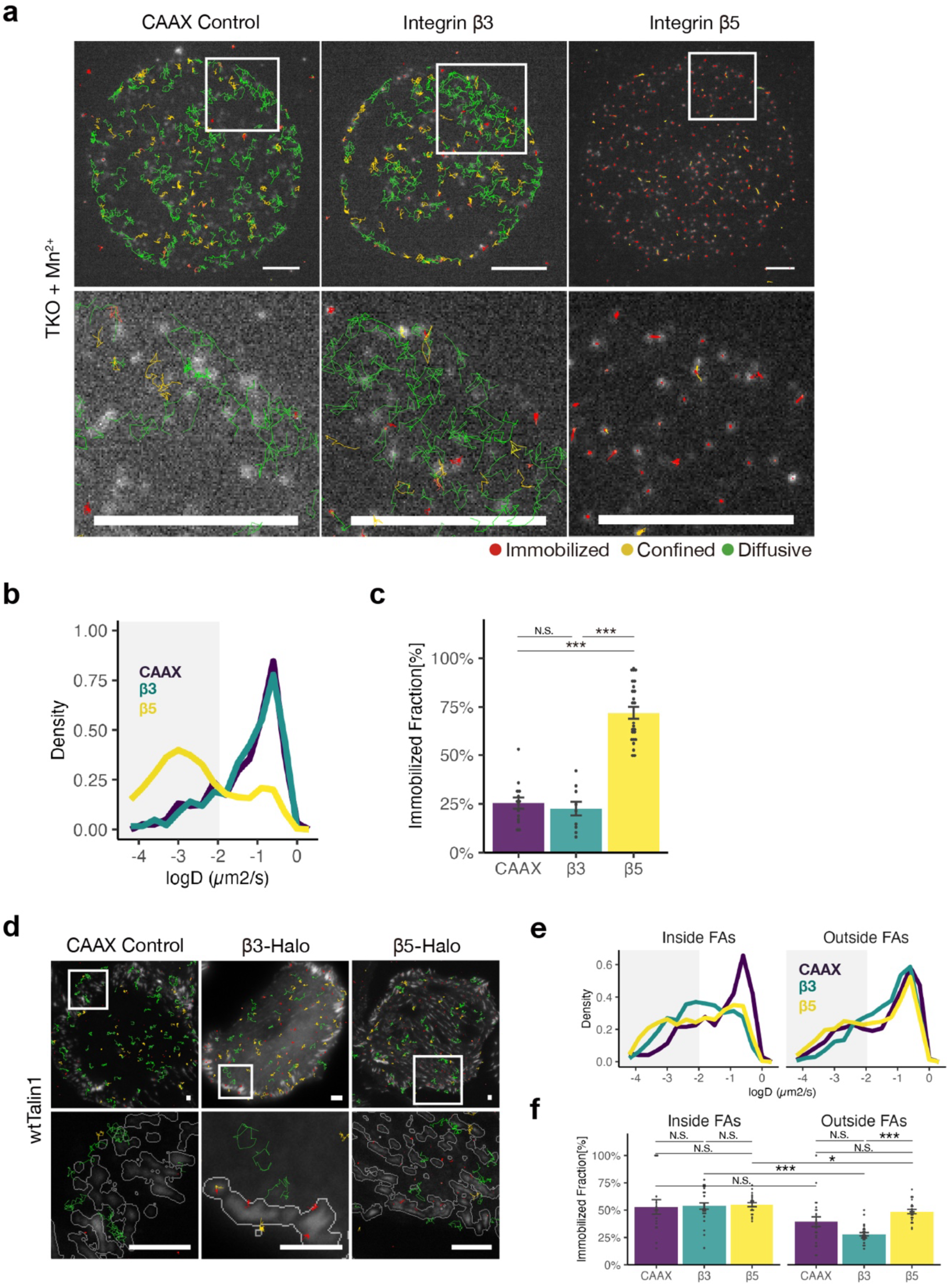
**Single-particle tracking of integrin proteins in the absence and presence of talin1. a–c**) TKO cells stably expressing Halo-CAAX, β3-Halo, or β5-Halo were replated into fibronectin-coated substrate in the presence of 0.2 mM MnCl_2_ to activate integrins. Images were captured for 100 frames at 50 ms of exposure (actual time interval 17 Hz) at least 4 h after replating. **a**) Single-molecule tracks of Halo-CAAX (left; **Movie 11**), β3-Halo (middle; **Movie 12**), and β5-Halo (right; **Movie 13**). Tracks are shown overlaid with single-molecule images. **b**) Distributions of the diffusion coefficient *D* computed from the tracks of Halo-CAAX (purple), β3-Halo (green), and β5-Halo (yellow). **c**) Fractions of Halo-CAAX (purple), β3-Halo (green), β5-Halo (yellow), undergoing immobilization (*D*<0.01). (**d–f**) TKO cells stably expressing wtTalin1-mEm and either Halo-CAAX, β3-Halo, or β5-Halo were replated and imaged. **d**) Single-molecule tracks of Halo-CAAX (left; **Movie 14**), β3-Halo (middle; **Movie 15**), and β5-Halo (right; **Movie 16**), co-expressed with wtTalin1-mEm. Tracks are shown overlaid with wtTalin1-mEm images and focal adhesion (FA) ROIs (insets). **e**) Distributions of the diffusion coefficient *D* computed from the tracks of Halo-CAAX (purple), β3-Halo (green), and β5-Halo (yellow), inside (left) and outside (right) of FAs. **f**) Fractions of Halo-CAAX (purple), β3-Halo (green), β5-Halo (yellow), undergoing immobilization (*D*<0.01) inside (left) and outside (right) of FAs. Sample sizes are as follows: (**a–c**) 563, 644, 2288 tracks from 14, 10, 25 cells, pooled from 3, 3, 3 experiments. (**d–f**) 365, 1350, 2847, 1061, 2296, 6861 tracks from 22, 27, 22 cells, pooled from 3, 3, 3 experiments. Statistical significances were obtained using Kruskal-Wallis rank sum test followed by pairwise Wilcoxon rank sum test. Bonferroni *P*-value adjustment was performed for multiple comparisons within either inside or outside FAs independently. N.S., no significance, \**P*<0.05, \*\**P*<0.01, \*\*\**P*<0.001. Error bars show mean ± s.e.m. Scale bars, 2 µm.

The single-construct nature of optodimerizable-talin1 enables us to generate TKO cells with stable expression of ΔDD-Dronpa together with another construct, such as the HaloTag fusion of CAAX membrane probes or integrin. SPT analysis was performed in these cells using brief illuminations of mIFP to locate FA regions for segmentation at the beginning and the end of the time-lapse series (Fig. 7a). As shown in Fig. 7b**–g**, diffusion coefficients corresponding to regions within FAs and outside of FAs were analyzed. CAAX mobility was similar between inside and outside FAs as expected (Fig. 7b**–d**). We observed that immobilized integrin β5 is relatively enriched in FAs, compared to outside of FAs, exhibiting similar behavior to in talin1-rescued TKO cells (Fig. 7h**–j**). In contrast, we observed that β3 was predominantly mobile in ΔDD-Dronpa for both fractions in FA and outside of FAs in contrast to wt-Talin1, and it was not affected by monomerization or subsequent re-dimerization (Fig. 7e**–g**). These results suggest that integrin β3 immobilization could be more sensitive to ABS3 actin-binding activity of talin, which may be somewhat attenuated in ΔDD-Dronpa as seen from actin retrograde flow engagement (Fig. 4a**,b**). Further investigations will be necessary to dissect the underlying mechanisms.

**Figure 7:**
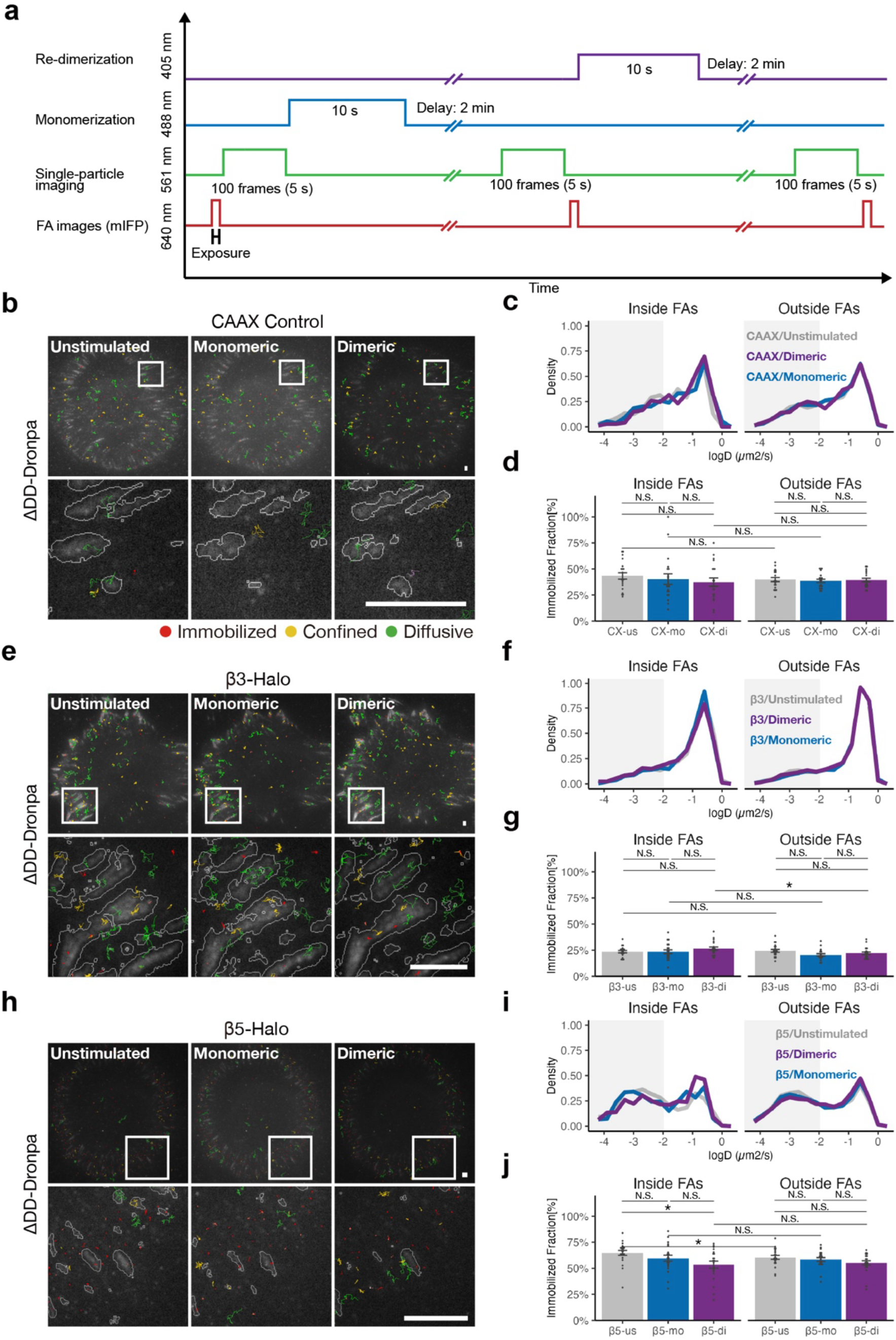
Multiplexing optogenetic control of talin dimerization and single-particle tracking of integrin proteins. **a**) Schematic diagram of optogenetic stimulation and image acquisitions. TKO cells stably expressing ΔDD-Dronpa, and either Halo-CAAX, β3-Halo, or β5-Halo were replated into fibronectin-coated substrate. Images were captured for 100 frames at 50 ms of exposure (actual time interval 17 Hz) at least 4 h after replating. **b,e,h**) Single-molecule tracks of Halo-CAAX (**b, Movie 17**), β3-Halo (**e, Movie 18**), and β5-Halo (**h, Movie 19**), co-expressed with ΔDD-Dronpa. Tracks are shown overlaid with mIFP channel of ΔDD-Dronpa images and FA ROIs (insets). **c,f,i**) Distributions of D computed from the tracks of Halo-CAAX (**c**), β3-Halo (**f**), and β5-Halo (**i**), in unstimulated state (gray), 2 min after monomerization (blue), and 2 min after re-dimerization following the monomerization (purple), respectively, inside (left) and outside (right) FAs. **d,g,j**) Fractions of Halo-CAAX (**d**), β3-Halo (**g**), and β5-Halo (**j**), in unstimulated state (left), 2 min after monomerization (middle), and 2 min after re-dimerization following the monomerization (right), respectively, inside (left) and outside (right) FAs. Sample sizes are as follows: (**c,d**) 2072, 2389, 1835, 4405, 3993, 4622 tracks from 20 cells, pooled from 3 experiments. (**f,g**) 606, 1215, 572, 4216, 4783, 4696 tracks from 21 cells, pooled from 3 experiments. (**i,j**) 264, 356, 193, 1914, 1862, 1896 tracks from 20 cells, pooled from 3 experiments. Statistical significances were obtained using Kruskal-Wallis rank sum test followed by pairwise Wilcoxon rank sum test. Bonferroni *P*-value adjustment was performed for multiple comparisons within either inside or outside FAs independently. N.S., no significance, \**P*<0.05, \*\**P*<0.01, \*\*\**P*<0.001. Error bars show mean ± s.e.m. Scale bars, 2 µm.

We next made use of the optodimerizable-talin1 to investigate how talin monomerization may affect the mobility of integrin β5. Monomerization of talin was performed using 488-nm illumination (2.8 mW, 10 s), while re-dimerization of talin was performed using 405-nm illumination (3.1 mW, 10 s; Fig. 7a). SPT tracking was performed subsequently by imaging the 561-nm channel as above. Control experiments using the CAAX probe in TKO cells stably expressing ΔDD-Dronpa, revealed that highly mobile CAAX trajectories predominate both inside and outside FAs (Fig. 7b). The diffusion coefficient distribution of CAAX appears minimally perturbed (Fig. 7c**,d**) upon 488-nm and 405-nm illumination to monomerize or re-dimerize talin, as expected. In contrast, a noticeable redistribution in integrin β5 mobility was observed, whereby upon 488-nm illumination, the immobilized fraction of integrin β5 decreased by 5 and 13%, 2 and 5 min after stimulation, respectively (Fig. 7i**, Fig. S5**). This suggests that talin dimeric state is important for the immobilization of integrin β5 in FAs, implying that FA-resident population of integrin β5 is probably dependent on talin1. Our observation also suggested that the monomerization of talin1 may liberate integrin β5 to de-adhere and diffuse away. Upon using 405-nm illumination for 10 s to activate talin dimerization, we observe that integrin β5 immobilization remains depleted, however. Re-dimerization by 405-nm light did not fully recover the immobilization of β5, at least within the time period we tested (10 min; Fig. 7g**–j****, Fig. S5**). This observed difference in β5 mobility profiles was confirmed to not be an artifact derived from photobleaching or random fluctuations (**Fig. S6**). Altogether, these results underscore the suitability of optodimerizable-talin1 in terms of multiplexing optogenetics with single-molecule-based super-resolution microscopy. This also opens up the possibility for future investigation in FA dynamics. For example, how different modes of integrin β5 activation and de-activation in FA and RAs could be regulated.

## Discussion

This study highlights the importance of dimerization in the function of talin within FAs and harness such effect to develop a molecular tool for mechanobiological mechanistic dissection. Using site-specific replacement of the native dimerization domain, DD, with an optically modulated dimerizer moiety, pdDronpa1.2, we showed that talin dimerization is an indispensable molecular event for FA formation and maintenance, as well as engagement with the actin retrograde flow and integrin β5 immobilization. The key feature of the Dronpa-based system we utilized in this study is its single-construct design. This is in contrast to previously reported optogenetic tools to control force transmission at FAs (Aureille et al., 2024; Sadhanasatish et al., 2023; Yu et al., 2020) which require the expression of two components, whose expression at the same level in the same cells is often challenging. Such stoichiometric challenge is even more acute for multiplexing experiments, where the expression of additional genetically-encoded probes on top of the optogenetic probes is needed. With the single-construct design, this problem can be significantly alleviated. For example, reporters of localization, conformation, or activity of FA-associated molecules, or mutants thereof, can be readily co-expressed to study their responses to, or effects upon, talin monomerization. This is demonstrated here with our actin retrograde flow analysis in different molecular clutch engagement regime achieved by light-based modulation of talin dimeric state, where distinct ‘molecular clutch’ behavior can be directly visualized. While in our study stable expression of optodimerizable-talin1 was applied, the single-construct design will also facilitate other approaches such as CRISPR/Cas9-mediated knock-in, allowing optogenetically-modified protein to be expressed at endogenous level. Notably, dimerization has been shown to be important for essential integrin-associated proteins such as kindlin and filamin (Liu et al., 2015a; Sun et al., 2019a), and thus they could potentially be optogenetically controlled by a similar single-construct design as well.

One limitation of our current optodimerizable-talin1 is that some functional aspects of wild-type Talin1 appears to be attenuated. In particular, while the expression of ΔDD-Dronpa in TKO cells induced cell spreading, polarization, FA formation and maturation to a comparable level as wtTalin1-expressing cells, actin retrograde flow engagement was only partially rescued. The rescue of FA formation by ΔDD-Dronpa indicate the ability to at least activate integrin β1 and β5, but differential effects on different integrins were observed with integrin β3 immobilization seemingly deficient in FAs within the observed timescale. These observed differences may arise from a number of reasons. For example, leakage of monomerization could arise due to the dimerization affinity for pdDronpa1.2 homodimer (K_d_ = 4 µM) (Jöhr et al., 2019) being somewhat weaker than that of other optically controlled heterodimerizer modules such as LOV2 and Zdk2 (K_d_ = 17 nM) of the LOVTRAP system, recently used to control talin (Sadhanasatish et al., 2023). In addition, recent Cryo-EM analysis of the ABS3 and DD domain revealed the participation of the DD domain itself in actin binding (Biertümpfel et al., 2025). Here, the ΔDD-Dronpa addition could perturb the nearby C-terminal integrin binding site, IBS2 (residues 1984–2344) which has been shown to engage in integrin β3 binding (Moes et al., 2007; Rodius et al., 2008). Thus, it is conceivable that the perturbation due to Dronpa replacement, together with monomerization leakage, may account for the attenuated molecular clutch behavior observed here. Nevertheless, new structural information as well as advances in artificial intelligence-based protein design (Kyro et al., 2025) would be particularly useful for improving the iteration of the optogenetic design.

By dissecting the C-terminal regions of talin, our study provides additional clues to the function of this domain. We previously used both chemogenetics and optogenetics approaches to modulate the cellular function of the ABS3 domain in talin, showing that it acts as a key regulator of actin and FA dynamics (Wang et al., 2019; Yu et al., 2020). This is in good agreement with previous evidence that ABS3 is vital for talin activation inside focal adhesions (Atherton et al., 2015; Rahikainen et al., 2019), although this can be bypassed by increasing the levels of active vinculin in the cell. By deleting the dimerization domain, or by using light-mediated dissociation, we have shown that monomeric talin is associated with adhesion degradation and increased actin flow rates. Together with recent Cryo-EM structural information (Biertümpfel et al., 2025), our observations suggest a model for talin engagement with rearward flowing actin, in which the dimerized talin may be ‘riding’ on top of an actin filament with the dimerization domain forming the apex of the contact with actin, while the R13 can be involved in providing friction through additional interactions with the actin filament allowing the generation of tension through talin that opens up additional actin crosslinking sites. Since actin filaments in the lamellipodia are known to comprise a substantial portion of dendritic meshworks, DD could potentially sterically engage, i.e. ‘catch’, at these branched points, generating tension in talin that is in turn transmitted to integrin and the ECM. Such engagement could persist for the lifetime of the dimer, but can also likely reform rapidly once disrupted due to the close proximity. Notably, purified talin has been characterized in both dimeric (Goult et al., 2013a) and monomeric autoinhibited form (Dedden et al., 2019). It is conceivable that both forms (and variants) exist in the cytoplasmic environment, especially since talin can undergo calpain cleavage to remove the dimerization domain (Bate et al., 2012). Talin dimerization was recently confirmed to be essential *in vivo*, as loss of dimerization abolished the synergistic interaction between talin ABS2 and ABS3 mutant alleles in Drosophila (Camp et al., 2024). However, what the cellular functions of talin in the monomerized form are is less clear, but may well involve shuttling to different cellular compartments beyond FAs, as increasingly observed in other adhesion proteins (Haage and Dhasarathy, 2023).

By multiplexing optogenetic control of talin dimerization with SPT analysis of integrin motion, this permitted dissection of factors regulating integrin mobility. We observed that talin monomerization resulted in a more diffusive behavior of integrin β5. This is in line with the previous reports showing tension-dependent localization of β5 into FAs (Zuidema et al., 2022; Zuidema et al., 2018). However, over a time period of several to ten minutes, re-normalization of talin does not appear to increase integrin β5 immobilization in FAs. Previous analysis of ultra-long trajectories has shown that integrin β1 and β3 undergo multiple rounds of immobilization and diffusion (Tsunoyama et al., 2018). It remains unclear whether integrin β5 is capable of similar dynamics. From our control experiments in TKO cells, in the absence of talin but with Mn^2+^ integrin β3 appears to be largely mobile while a significant fraction of integrin β5 is immobile, suggesting that integrin β3 could be more sensitive to talin-mediated engagement than β5. Notably, beyond its FA localization, integrin β5 is also known as a key component of reticular adhesions (RAs), which are dependent on clathrin-mediated endocytosis and are regulated by distinct signaling pathways (Lock et al., 2018). β5 has been shown to relocate from FAs into RAs upon S759/762 phosphorylation, while interconversion of FAs and RAs can occur via selective exchange of components on a stable αVβ5 scaffold (Zuidema et al., 2022). One possibility is that molecular signaling events i.e. phosphorylation may constitute additional molecular factors beyond talin dimerization required for integrin β5 immobilization. This may enable integrin β5 to act as a mechanosignalling integrator whose activation depends on both talin-mediated mechanobiological cues with biochemical signaling. Future detailed analysis will be necessary to elucidate such processes.

In summary, in this study we demonstrate how light-based control of force transmission at FAs can be engineered using a single-construct approach that streamlines experimental procedure and offers multiplexing capability enabling adhesion control in conjunction with advanced imaging methods. Our study also helps delineate the overlapping roles of the C-terminal actin-binding/dimerization domain of talin, showing that the actual dimerization process itself contribute significantly to the formation and maintenance of FAs. Distinct molecular behaviors of different integrin heterodimers were unveiled, highlighting the utility of our approach. Further development of this strategy should enable systematic molecular dissection of mechanobiological processes with spatiotemporal precision.

## Methods

### DNA constructs and molecular cloning

Fluorescent proteins (FP) fusion constructs were based upon either N1-or C1-Clontech-style vectors with CMV promoters (from Dr. Michael Davidson, The Florida State University, Tallahassee, USA). For mApple-talin1(1–1973)-pdDronpa1.2, the construct contains the mApple fluorescent protein at the N-terminus, conjugated with mouse talin1 (residue:1−1973) and pdDronpa1.2 at the C-terminus. For pdDronpa1.2-talin1(1974–2541)-mIFP, the construct contains pdDronpa1.2 at the N-terminus, conjugated with mouse talin1 (residue: 1974−2541), followed by mIFP fluorescent protein at the C-terminus. For ΔDD, the construct contains the mIFP at the N-terminus, conjugated with mouse talin1 (residue: 1–2493). For ΔDD-Dronpa, the construct is based on the ΔDD construct and followed by mIFP at the C-terminus. For the β3-Halo, HaloTag sequence (Promega) was PCR amplified and cloned into the Integrin-beta3-PAGFP plasmid (Changede et al., 2019) by replacing the PAGFP. For the Halo-CAAX plasmid, the CAAX sequence from the human H-RAS transcript and the HaloTag sequence were cloned into the EGFP-N1 plasmid by replacing the EGFP. For the β5-Halo, (SGGGG) _3_-Halo7 sequence from pEGFP-N1/Integrin-b3-SGx3-Halo7 plasmid (from Dr. Akihiro Kusumi, Okinawa Institute of Science and Technology, Japan) was cloned into the Integrin beta5-2xEGFP plasmid (from Dr. Staffan Strömblad; Addgene plasmid #139779) by replacing EGFP, and then Q165H/P174R point mutations were introduced to mutate Halo7 into Halo9. Molecular cloning above was performed by either internal or external services (Epoch Life Sciences, USA), verified by whole-plasmid sequencing, and amplified by QIAGEN Plasmid Midi Kit (QIAGEN, USA). The following constructs are generous gifts from external sources: mCh-wtTalin, wtTalin-mEm, TalinH, and mCherry-Vinculin are from Dr. Michael Davidson (The Florida State University, Tallahassee, USA) and F-tractin-mCherry is from Dr. Michael Sheetz (Mechanobiology Institute, Singapore) respectively. More details on other constructs can be found in **Table 1**.

Expression vectors were generated by gene synthesis (Epoch Life Sciences, USA), verified by sequencing, and amplified by QIAGEN Plasmid Midi Kit (QIAGEN, USA).

### Cell culture and transfection

Mouse kidney fibroblasts lacking both Talin1 and Talin2 (TKO cells) have been described previously (Austen et al., 2015) and were generously gifted by Prof. Carsten Grashoff (University of Münster, Germany). TKO cells were maintained in Dulbecco’s Modified Eagle’s Media (DMEM) with GlutaMAX supplemented with 10% FBS, 1% Sodium pyruvate, and 1% Penicillin/Streptomycin cocktail and cultured in a 37°C incubator with 5% CO_2_. For transfection, cells were diluted to a density of 6 × 10^6^ cells/mL and mixed with 0.5−2.0 μg of expression vectors per each electroporation reaction. Transfection was performed by Neon Electroporation System (Thermo Fisher) per the manufacturer’s protocol, with the settings of 1,100 V, 30 ms, 2 pulses at 10-µL scale. For imaging, transfected cells were sparsely plated at a density of 7000/cm^2^ on fibronectin-coated coverslips or glass-bottom dishes (Iwaki, Japan). Only clearly transfected cells were measured and imaged. Cells were screened monthly for any mycoplasma contamination.

### Live Cell Microscopy and Optogenetic Manipulation

TIRF imaging of live cells was performed using a Nikon Eclipse Ti inverted microscope (Nikon Instruments), equipped with a motorized TIRF/FRAP illuminator (iLas2, Gataca Systems), with a polarization-maintaining optical fiber-coupled laser combiner (100 mW 405 nm, 60 mW 488 nm, 50 mW 561 nm, and 100 mW 642 nm solid-state lasers, Omicron Laserage), a light emitting diode-based epifluorescence excitation source (SOLA, Lumencor), an ORCA-flash 4.0 sCMOS camera (Hamamatsu), a 100× N.A. 1.49 Apo TIRF objective lens (Nikon Instruments), and an Okolab stage-top chamber with CO_2_ and temperature control (Okolab, Italy). Activation (dimerization) and deactivation (monomerization) of the pdDronpa system are performed with either 405-or 488-nm laser in the widefield configuration.

### Multiplexed Optogenetic control and Single-Molecule imaging

Multiplexed optogenetic control and single-molecule imaging were performed using the setup described below. For visualization of single Halo-tagged molecules, cells were harvested, incubated at 37°C for 30 min in suspension, plated on glass-bottom dishes, and incubated at 37°C for 3 h. They were then treated with either 0.1 nM Janelia Fluor 549 or 646 HaloTag Ligand in culture medium for 15 min, 37°C, rinsed twice with imaging medium [FluoroBrite DMEM (Thermo Fisher) supplemented with GlutaMAX, 10% FBS, 1% sodium pyruvate, and 1% Penicillin/Streptomycin cocktail] and then replaced with fresh imaging medium. All the samples were observed at least 4 h after replating. Light stimulations with either 405-or 488-nm lasers were conducted at 7.5% of the maximum power for 10 s continuously without ROI. Single-molecule image stacks were recorded for 100 frames at 50 ms of exposure (58 ms of actual time interval) before stimulations, and after each stimulation (by 488-and then 405-nm light) with 2–10 min of delays. Images of the mIFP channel visualizing ΔDD-Dronpa were also captured at the beginning (for the “Unstimulated” condition) and after each acquisition of single-molecule image stacks (for the “Monomeric” and “Dimeric” conditions), which were used for generation of FA mask for track classification.

### Single-molecule Tracking and analysis

Custom Python software was developed for single-molecule tracking and analysis. For single-molecule detection, raw camera frames were first converted into photon unit (*N*) using the quantum efficiency curve provided by the manufacturer, and the equation:

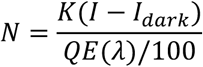

where *K* is the conversion factor; *I*, the pixel intensity; *I_dark_*, the dark offset; and QE(λ), the quantum efficiency at the specified wavelength λ. Single-molecule detection was performed using a previously described algorithm (Kanchanawong and Waterman, 2013; Shtengel et al., 2014). Briefly, Laplacian of Gaussian filter with radius of 2 pixels was used to enhance point spread function (PSF)-like objects to identify single-molecule peaks. Peak candidates were first identified by performing a local maxima search with a 5×5 pixels moving window, ranked by intensity, and subjected to cut-off based on minimum photon number criteria. High-precision xy-coordinate of each molecule is obtained by localization analysis using the Gpufit library on NVIDIA CUDA graphics card (Przybylski et al., 2017), using the asymmetric 2D Gaussian fit model. The coordinates obtained were then linked using the trackpy library (Crocker and Grier, 1996) with a search radius of 8 pixels (853.36 nm) and a gap parameter of 0 frame. Sufficiently long trajectories with at least 25 frames were chosen for further analysis. The diffusion coefficient for each trajectory (*D*) was calculated by performing a linear fitting to the first 4 datapoints of mean-square-displacement (MSD) vs lag-time, *t,* as calculated by trackpy:

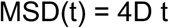

The degree of confinement, r_conf_, for each trajectory was calculated as described earlier (Rossier et al., 2012), where MSD(t) was fitted to:

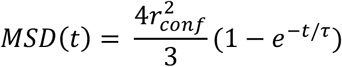

where r_conf_ is the confinement radius. Additionally, to improve robustness in trajectory classification, the moment-scaling analysis (Ewers et al., 2005; Ferrari et al., 2001) was performed by calculating the moment 0,1,2,..,6 of displacement, μ^v^, as defined by:

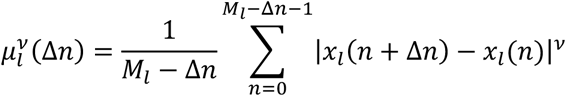

where *x_l_(n)* is the coordinate vector at time *nΔt*; *Δn*, the specific frame shift; *M_l_*, the maximum number of points in trajectory *l* used for the analysis. Subsequently, the scaling coefficients γ^v^ for each moment (1…6) are calculated by linear regression of *log μ^v^* versus *log nΔt.* Finally, the moment scaling spectrum (S_MSS_) is calculated as the slope of *γ^v^* versus *v* plot. For strongly self-similar processes, S_MSS_ is considered to provide a robust indicator of motion type, whereby S_MSS_ corresponds to 0 for stationary particles, ½ for freely diffusing particles, and 1 for directed motion. S_MSS_ calculations were performed using the lag time fitting up to ¼ of the total points in the trajectory. Segmentation of the focal adhesion region was performed with ilastik (Berg et al., 2019) on mEmerald (wtTalin1-mEm) and mIFP (ΔDD-Dronpa) fluorescence channels, respectively, followed by 1 px of dilation on binary images by ImageJ. Trajectories were classified to be in a region-of-interest (ROI) if the center-of-mask of the coordinates is within the ROI.

### Immunofluorescence microscopy

Cells were fixed in 4% formaldehyde in PEM buffer (80 mM PIPES, 1 mM EGTA, 1 mM MgCl_2_, pH 6.8) for 10 min at 37°C and then permeabilized with 0.2% triton X-100 in phosphate buffer. Samples were then blocked with 4% BSA in phosphate buffer for one hour and stained with antibody overnight at 4°C. Primary antibodies and dilutions were: pY397-FAK 1:300 (Abcam ab81298), Vinculin 1:200 (Sigma-Aldrich V9131), pY118-Paxillin 1:50 (Cell Signaling 2541S), active integrinβ1 1:200 (BD 553715). Secondary antibody staining was with either Alexa Fluor 568-conjugated Goat anti-Rat 1:200 or Alexa Fluor 647 conjugated Donkey anti-Rabbit, Donkey anti-Mouse, or Goat anti-Rat antibodies 1:50 (Thermo Fisher A11077, A31573, A31571, A21247, respectively). F-actin was visualized with phalloidin-CF405 (Biotium BT00034-T) at 1:50 dilution.

### Spinning disc confocal microscopy

Confocal imaging was performed using a CSU-W1 Spinning Disk (Yokogawa) confocal microscope with a single, 70-μm pinhole disk and quad dichroic mirror along with an electron-multiplying charge-coupled device (EMCCD) (Princeton Instruments, ProEM HS 1024BX3 megapixels with 30-MHz cascade and eXcelon3 coating) using a Plan Apo 100× NA 1.45 objective lens.

### YAP Quantification

YAP nuclear to cytoplasmic ratio was calculated in a similar way as reported before (Elosegui-Artola et al., 2017). YAP1 (Abnova H00010413-M01; 1:200) was immunostained and imaged using a spinning disc confocal microscope with 0.3 µm z-step size. The plane of the maximum nuclear area was assessed by co-immunostaining with DAPI (Invitrogen D3571; 1:200). A 10×10 pixel square region was drawn inside the nucleus, and an equal-sized region was drawn in the cytosol immediately adjacent to the nuclear region. YAP nuc/cyt ratio was calculated from YAP fluorescence intensities measured in both these regions after background subtractions.

### Analysis of actin retrograde flow

TKO cells were transfected with either Talin1/ΔDD/ΔDD-Dronpa in combination with F-tractin-mCherry or F-tractin-mCherry alone and spread with 2 mM MnCl_2_ which activates integrins. The cells were spread on fibronectin-coated dishes (Iwaki) and imaged on a Nikon Ti-E equipped with a W1 spinning-disk confocal and LiveSR super-resolution module (Roper Scientific). Images were acquired with a Nikon 100x TIRF 1.49 NA lens onto a Prime 95B sCMOS camera (Photometrics). The acquisition was set at a 2-second interval for two minutes to capture actin flow rates in the lamellipodia. Kymographs were produced using the kymograph plugin available in FIJI, and PIV analysis was performed using the matPIV toolbox for MatLab (Sveen, 2004). PIV maps were averaged over a 12-second period to reduce noise and characterize the actin flow rates more accurately. To compare the effects of dimeric and monomeric talin on actin flow, cells were prepared as described above, but with actin flow imaged at 1-second intervals over 5 seconds with 1 minute between sequences. After two five second series were collected to obtain the baseline, talin was monomerized with 488 nm light as monitored by the loss of pdDronpa1.2 fluorescence. Five image series were collected coincident with 488-nm illumination, and then this process was repeated with 405-nm light to redimerize talin.

## Statistical analysis

Data are expressed as mean ± standard error (SE) of at least three independent experiments. All statistical analyses were performed in either GraphPad Prism or R software. *P*-values were calculated using the Mann-Whitney U test for the comparison of two groups, and the Kruskal-Wallis rank sum test followed by the pairwise Wilcoxon rank sum test with Bonferroni *P*-value adjustment was performed for more than three groups, respectively. In all cases, *P*<0.05 was considered statistically significant.

## Code availability

Python codes for single-molecular tracking are available upon request.

## Data availability

All data supporting the findings of this study are available from the corresponding author upon reasonable request.

## Acknowledgement

The authors thank the microscopy, IT, high-throughput molecular genetics, and wet lab core facilities of the Mechanobiology Institute (MBI) for infrastructure support. We also thank Drs. Akihiro Kusumi and Takaaki Tsunoyama (Okinawa Institute of Science and Technology, Japan), and Dr. Cheng-Han Yu (The University of Hong Kong, Hong Kong SAR, China) for their helpful comments. The research is funded by Ministry of Education Singapore Academic Research Fund Tier 2 (MOE-T2-1-124, to P.K.), Ministry of Education Singapore Academic Research Fund Tier 3 (MOE-T3-2020-01, to P.K.), MBI intramural funding (P.K.), and National Research Foundation Singapore (NRF-MSG-2023-0001, to P.K.). K.J. is supported by MBI Graduate Fellowship. S.F.H.B. is partially supported by the National Research Foundation Singapore Quantum Engineering Programme (QEP-P-7).

---

**Supplementary Table 1:** Expression vectors used in this study.

| **Short form in text** | **Construct name** | **Source** |
| --- | --- | --- |
| mCh-wtTalin | mCherry-Talin1(1–2541) | Dr. Michael Davidson |
| wtTalin-mEm | Talin1(1–2541)-mEmerald | Dr. Michael Davidson |
| TalinH | EGFP-Talin1(1–433) | Dr. Michael Davidson |
| - | mApple-Talin1(1–1973)-pdDronpa1.2 | This work |
| - | pdDronpa1.2-Talin1(1974–2541)-mIFP | This work |
| ΔDD | mIFP-Talin1(1–2493) | This work |
| ΔDD-Dronpa | mIFP-Talin1(1–2493)-pdDronpa1.2 | This work |
| - | F-tractin-mCherry | Dr. Michael Sheetz |
| - | mCherry-Vinculin | Dr. Michael Davidson |
| Halo-CAAX | Halo7-CAAX | This work |
| Intβ3-Halo | Integrinβ3-Halo7 | This work |
| Intβ5-Halo | Integrinβ5-(SGGGG)<sub>3</sub>-Halo9 | This work |

